# Temperature is a key driver of a wildlife epidemic and future warming will increase impacts

**DOI:** 10.1101/272369

**Authors:** Stephen J. Price, William T.M. Leung, Chris Owen, Chris Sergeant, Andrew A. Cunningham, Francois Balloux, Trenton W.J. Garner, Richard A. Nichols

**Author notes:** These authors contributed equally to this work (Francois Balloux, Trenton W.J. Garner, Richard A. Nichols).

## Abstract

Increasing environmental temperatures are predicted to have increasingly severe and deleterious effects on biodiversity. For the most part, the impacts of a warming environment are presumed to be direct, however some predict increasingly severe disease epidemics, primarily from vector-borne pathogens, that will have the capacity to deplete host populations. Data to support this hypothesis are lacking. Here we describe increasing severity of ranavirosis driven by increasing temperature affecting a widely distributed amphibian host. Both *in vitro* and *in vivo* experiments showed that increasing environmental temperature leads to increased propagation of ranavirus and, in the latter, increased incidence of host infection and mortality. Also, temperature was shown to be a key determinant of disease dynamics in wild amphibians, raising the odds and severity of disease incidents. The direction of this effect was highly consistent in the context of other interacting variables such as shading around ponds. Projections based on future climate indicate that changes in seasonal weather in the UK will result in the increased incidence of severe cases of ranavirosis in amphibian populations that could affect recruitment. These complementary lines of evidence present a clear case of direct environmental modulation of a host-pathogen interaction and provide information for proposing mitigation actions.

---

The interaction between hosts, pathogens and their shared environment shapes infectious disease outcomes and the invasiveness of pathogens^1^. Climatic conditions at a landscape scale represent a critical dimension of the host environment but also directly affect pathogen survival and transmission^1–3^. As such, climate plays an important role in the rate and pattern of invasions by emerging pathogens and will help to predict emergences under future climate change scenarios. Unsurprisingly, most research effort in these fields has focused on human diseases^4^, with vector-borne diseases a target of particular research interest since these are expected to be affected to a greater degree by climatic variables^5^.

Human disease systems are restricted in their suitability for controlled experimental manipulation. Studies of ongoing human epidemics tend to be narrow in their approach, as medical intervention always takes precedence over more fundamental scientific questions, including the effect of temperature as a driver of disease outbreaks, and studies are further complicated by a series of cultural, economic or societal confounding factors^5,6^. Research on past epidemics, such as plague in the middle ages (e.g. Büntgen et al.^7^), suffers from poor quality epidemiological data and major difficulties in accessing biological material. Furthermore, the scope of concerning infectious disease issues far exceeds that affecting humans, and many non-human pathogens have the capacity to impact human well-being through their impacts on crops, livestock and wildlife^8,9^.

Increasing environmental temperature has been invoked as a key component of climate change which drives infectious disease emergence and severity, but it is often difficult to discriminate between the effect of temperature and other epidemiological determinants of disease outbreaks^10^. The direct and indirect influences of temperature on host-pathogen interactions^11^, its nonlinear effects on incidence and severity^12–14^ and its possible upstream effects^15,16^, all represent considerable challenges to a better understanding of disease emergence. However, research on non-human diseases can enable the complex effects of climate to be unpacked through rigorous evaluation and theoretical, observational, and experimental approaches. From this body of evidence, a robust understanding of the role of the environment in disease occurrence will arise which then enables projections of changes in risk^3,17,18^.

The growth of amphibian ranaviruses (large double-stranded DNA viruses; family *Iridoviridae*) is affected by temperature^19–21^ and this natural system has promising credentials as a model to explore interactions between pathogens, hosts and this aspect of the environment. The strength of the amphibian-ranavirus model is founded on a combination of: i) the availability of extensive epidemiological data from a long-term study of disease in wild populations of UK common frogs^22–24^, ii) the capacity to manipulate viral genetics and the pathogen’s environment - both host cell and growth conditions - in the laboratory, and iii) the possibility to replicate *in vitro* experiments *in vivo* under controlled conditions using natural hosts. In this study, we combined epidemiological modelling with all of the above to investigate the role of temperature as a driver of disease outbreaks in common frogs infected with ranaviruses in the frog virus 3 (FV3) lineage.

## Results and discussion

### Effect of temperature on disease incidence and severity in the wild

Ranavirus emerged in UK common frog (*Rana temporaria*) populations in the late 1980s following multiple introductions and continues to cause severe population declines^22,24–26^. Disease outbreaks are seasonal, peaking in summer months^23^, but the detectability of the main UK host, the common frog, is also strongly seasonal and there has been no previous attempt to explicitly control for host population density, host activity or observer effort in examining the periodicity of outbreaks. We used data from the Frog Mortality Project (FMP), a flagship citizen science project which has been collating reports of amphibian mortality incidents from members of the UK public for twenty-five years^24^. The dataset has been reliably filtered for incidents of ranavirosis^24,26,27^ and, in our study, the seasonal detectability of amphibians was controlled for indirectly through the inclusion of mortality incidents caused by factors other than ranavirosis as previously^24^. The finalized FMP dataset used in this study contains 4385 unique records, of which 1497 are classed as ranavirus-consistent.

A simple logistic regression model using this full dataset, and with temperature as the sole predictor of ranavirus status (Model 1), revealed a highly significant effect of temperature on the proportion of ranavirus-consistent incidents, reducing the deviance by 52.6 compared to the null model (Χ^2^ test; p = 4.02 × 10^-./^). For each 1°C increase in temperature, the odds that an incident was caused by ranavirus increased by 4.4% (p = 8.89 × 10^-./^). A model with a transition between an upper and lower frequency (Model 2) significantly improved the fit to the data compared to Model 1 (AIC scores were 5565 and 5581 respectively). Model 2 shows a step-change: below approximately 16°C, 25.1% of incidents were ranavirus-consistent, rising to 38.5% after the temperature threshold was crossed (Figure 1a). The difference between incidents that were ranavirus-consistent and the remainder (’non-ranavirus‘) is also apparent in the distribution of temperature records: the non-ranavirus category being strongly bimodal with peaks at both low and high temperature, whereas most of the ranavirus-consistent incidents were reported at higher temperature (Fig. S1).

**Figure 1.**
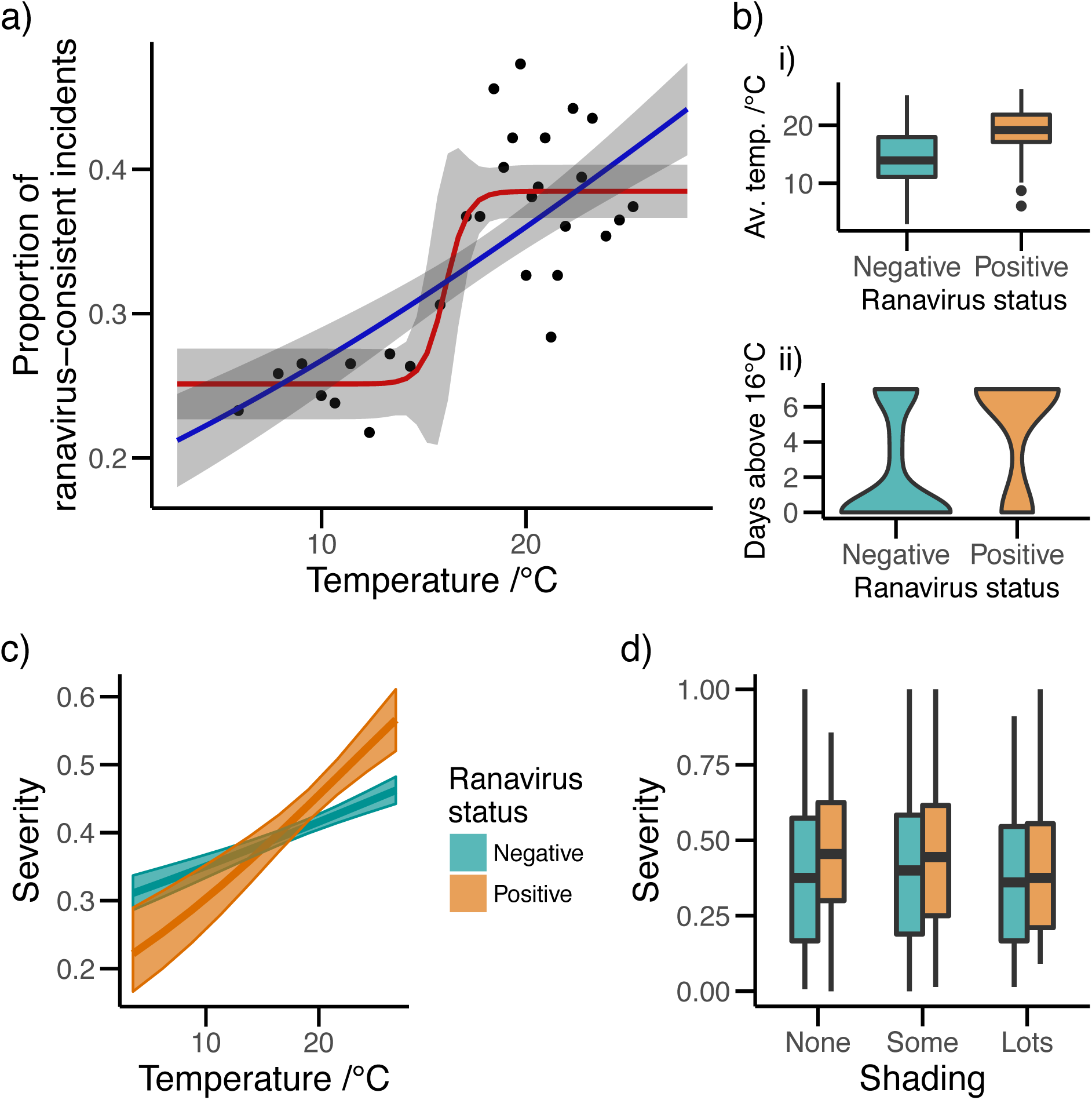
Warm temperatures increased the odds of occurrence and severity of incidents of ranavirosis involving wild common frog populations of the United Kingdom. a) The effect of temperature on the proportion of citizen science reports of frog mortality that were classified as ranavirus-consistent. The blue line is the fit from a simple logistic regression with the average maximum daily temperature for the month of onset of mortality as a predictor of ranavirus status. The red line is the maximum likelihood model of a logistic transition between a lower and upper frequency. Shaded areas around the lines represent 95% confidence intervals, calculated using the delta method for the red line. Points represent the observed data, grouped in windows containing equal numbers of records. b) Temperature in the week preceding frog mortality incidents confirmed by molecular methods predicted the ranavirus status. i) Average daily maximum temperature in the seven days preceding incidents by ranavirus status (“Positive” versus “Negative”). ii) The number of days in the week preceding mortality incidents where the daily maximum temperature exceeded 16°C by ranavirus status. c) The severity of frog mortality incidents (estimated proportion of population that died) was consistently greater at higher temperatures, particularly in the case of ranavirus-consistent incidents. The plot shows fitted lines (and 95% confidence intervals) from a generalised linear model (quasibinomial regression) of severity as a function of ranavirus status and temperature (average daily maximum temperature for the month of onset of mortality incidents). d) The presence of shading reduced the severity of ranavirus-consistent mortality incidents.

We previously screened a UK amphibian and reptile tissue archive and found 39 ranavirus-positive mortality incidents out of a total of 229 incidents^28^. Temperature was again a highly significant predictor of ranavirus status when these records with precise timestamps, and where ranavirus status had been confirmed, were analyzed. This more precise information about timing enabled a fine-scale examination of the effect of temperature in the days preceding incidents. The average temperature in the seven days preceding incidents was a significant predictor of ranavirus status (p = 1.18 × 10^-0^; residual deviance of model = 154.2 on 195 degrees of freedom; Fig. 1b.i), with each 1°C increase in temperature increasing the odds that incidents were caused by ranavirus by 20%. The temperature threshold where the proportion of ranavirus incidents increased sharply in the analysis of the full FMP dataset was approximately 16°C. A second model - with the number of consecutive days where the daily maximum temperature in the week preceding incidents exceeded 16°C as a predictor - also indicated that warmer temperatures were a good predictor of ranavirus status (p = 2.27 × 10^-1^; residual deviance of model = 151 on 195; Fig. 1b.ii): each additional warm day raised the odds that incidents were caused by ranavirus by 33%. The 16°C threshold model had a slightly lower AIC score than the model using the average temperature as a predictor (155 compared to 158).

The FMP database contains data on the severity of outbreaks (the estimated proportion of the frog population that died). After removing records with missing values, we produced a dataset for investigating severity that contained 2667 records, of which 427 incidents were classified as ranavirus-consistent. In a simple logistic model of severity with ranavirus status, average daily maximum temperature for the month of onset of mortality, and their interaction as predictors, all three terms were significant predictors and retained in the minimum adequate model. Temperature explained the most deviance with each 1°C increase in temperature leading to a 2.8% increase in the proportion of the population that died (p = 1.61 × 10^-2^). There was a significant interaction between the two main effects as a consequence of the different effect of ranavirus status on the relationship between temperature and the proportion dead (p = 0.001): at low temperatures, the severity of ranavirus-consistent incidents was slightly lower than for other types of incident but at higher temperatures it was the ranavirus-consistent incidents that were more severe (Fig. 1c). Attempting to incorporate time (the year that mortality incidents began) did not result in an extension of the minimum adequate model, however data on the number of surviving frogs were not collected after 2000 meaning that this analysis was restricted to the years 1991 to 2000 only.

The effects of other covariates previously identified as having an influence on the occurrence or severity of ranavirosis in UK common frogs^27^ were also explored using a more complex model containing ranavirus status, temperature, log-transformed pond volume, the interactions of the three, shading around ponds, the presence of toads, newts and fish, as well as the region. After model simplification, the minimum adequate model retained all three terms from the simple model but there were also significant effects of the presence of toads, the presence of fish, shading, pond volume plus its interaction with temperature, and region (Table S1). Toads reduced the severity of mortality incidents whilst the presence of fish increased severity (Fig. S3) as found previously^27^. Shading decreased the severity of incidents (Fig. 1d). Notwithstanding, and irrespective of which covariate was considered, the effect of increasing temperature increased the severity of disease. This is perhaps best illustrated by the effects of pond shading, where increasing the amount of shading (and, presumably, decreasing the maximum temperatures that frogs would have been exposed to) was associated with reduced severity of ranavirosis and a decreasing disparity in the severity of incidents between ranavirus-consistent and non-ranavirus incidents (Fig. 1d).

### *In vitro* assessment of viral growth rates and *in vivo* tests of virulence

We examined *in vitro* viral growth rates using two UK isolates of FV3 (RUK11 and RUK13; see methods for detailed descriptions of isolates) and two cell lines. Each isolate was incubated at a range of temperatures up to 30°C with each cell line, but regardless of cell line or isolate, increasing temperature resulted in exponentially increased growth rates and rates of plaque formation (Fig. 2). A linear model of log viral titres against temperature, cell line and their interaction revealed significant effects of temperature (coefficient = 1.23, p < 0.001) and host cell line (IgH2 compared to EPC, coef = -8.64, p < 0.001) but no interaction (analysis of variance, comparing model with interaction term to a model with main effects only: Fdf = 1 = 0.0047, p = 0.95), indicating the overall effect of temperature was independent of the host environment.

**Figure 2.**
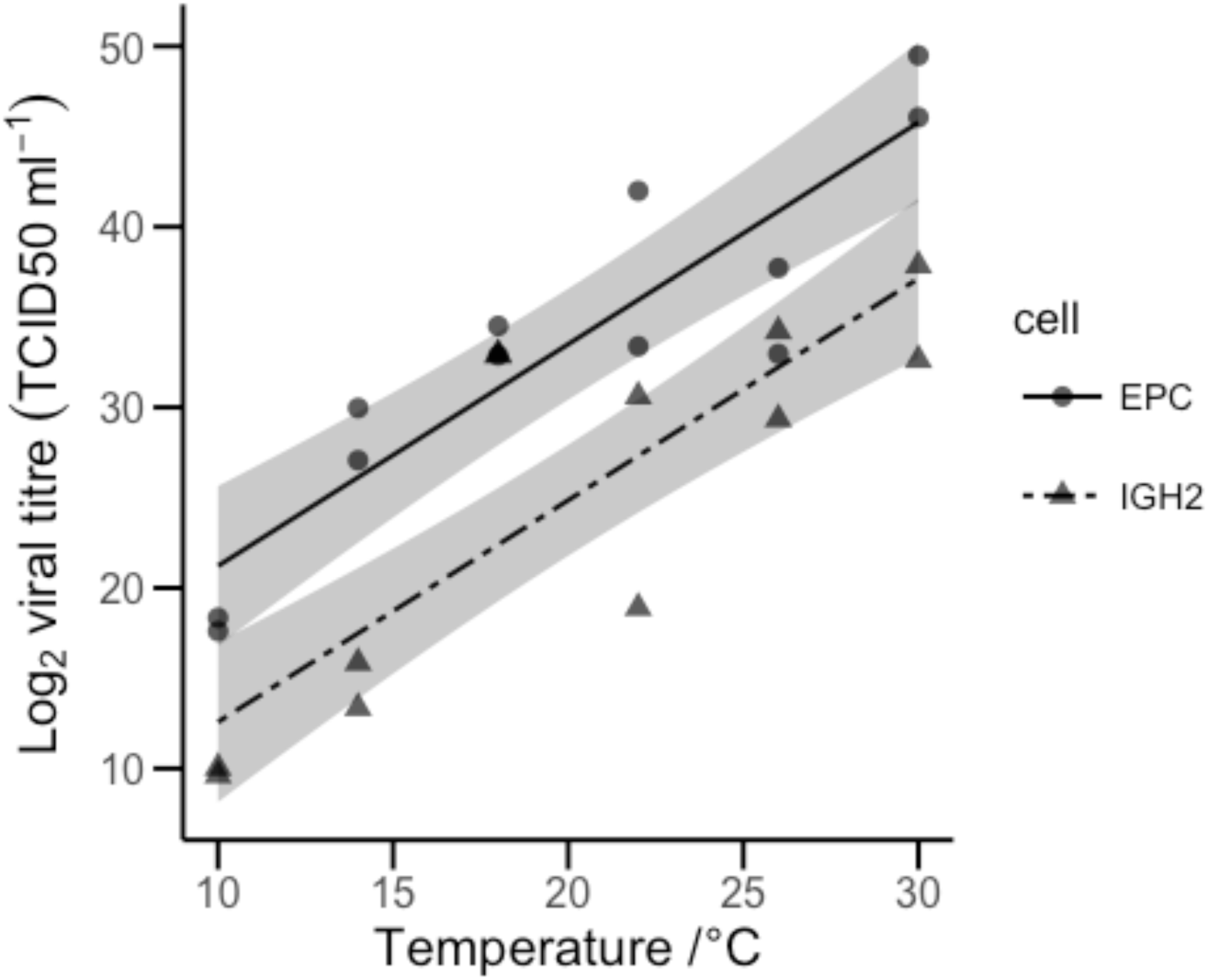
Effect of environmental temperature on growth of UK *Frog virus 3* (FV3) *in vitro*. Observed data (points) and predictions from linear model (lines with 95% confidence interval shaded) of FV3 growth at a range of environmental temperatures in fish (EPC; solid line) and reptile (IgH2; dashed line) cells. Growth was measured using the TCID50 method and is shown on a log scale. An increase in temperature of 1°C results in more than a doubling of viral growth (2.34 times).

In order to validate results from cell culture models *in vivo*, 60 overwintered common frog metamorphs (*R. temporaria*) were randomly allocated to one of six treatments (10 animals per treatment): three exposure treatments (sham, RUK11, RUK13) crossed with two temperatures (20°C [“low”] and 27°C [“high”]). Temperature was a highly significant predictor of survival: 20 of 60 animals died or were euthanized on reaching humane endpoints, of which 14 were from high temperature treatments and six were from low temperature treatments (Fig. 3a). Overall there was a 5.33 times higher risk of death in the high temperature treatments (p = 0.005; Fig. 3b). Titres of viral inoculates were not equalised between isolates. All individuals exposed to RUK11 (at a high dose) and maintained at high temperature died or reached endpoint by the eighth day post-exposure compared to six of ten individuals maintained at low temperature (Fig. 3a). Of the animals exposed to RUK13 (at the relatively low dose compared to RUK11), three individuals died or reached endpoint in the high temperature treatment compared with none at the low temperature. These results are largely in line with a previous study examining survival of common frog tadpoles exposed to a North American isolate of FV3, which showed that mortality was increased at 20°C compared to 15°C^19^. There was also a significant effect of exposure treatment: the expected hazard of animals exposed to RUK11 was 41.6 times higher than animals receiving a sham exposure (p = 0.0004). The expected hazard of animals exposed to RUK13 was only 3.23 times higher and was not significant (p = 0.31; Fig. 3b), but this was likely explained by the different doses used since both isolates had a similar effect on survival previously when titres were equalised^29^.

**Figure 3.**
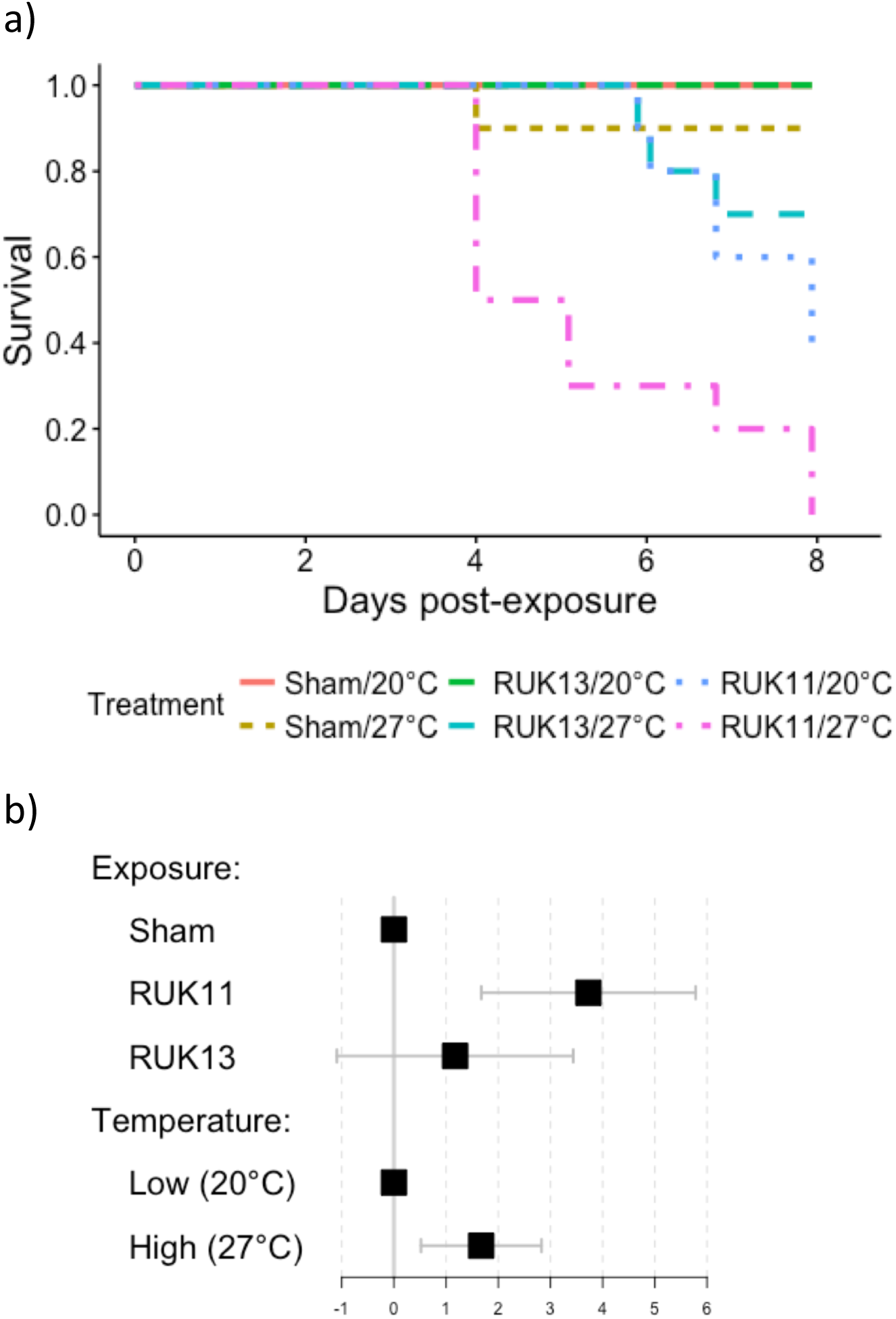
Effect of temperature and FV3 exposure on survival of common frogs. Survival was measured in each of six treatments; three exposure treatments (Sham, RUK11, RUK13) at each of two temperatures (20°C [“low”] or 27°C [“high”]). a) Kaplan-Meier survival plot. b) Forest plot of coefficients (± standard error) from a mixed effects Cox Proportional Hazards model of survival in response to exposure and temperature treatments.

Temperature was not retained as a significant predictor of *in vivo* viral load when logarithms of viral loads were modelled as a function of exposure and temperature treatments, survival outcome, and the interactions. However, the minimum adequate model did retain exposure (F251 = 277, p < 0.001), survival outcome (F1, 51 = 197, p < 0.001), and their interaction (F2, 51 = 25.7, p = 1.88 × 10^-4^; Fig. S4). Background levels of virus were detected from tissues of some animals subjected to a sham exposure but average viral loads were significantly higher in both frogs exposed to RUK13 (loads were 3.11 × 10^4^ times higher on average; p = 2.47 × 10^-5^) and frogs exposed to RUK11 (loads 1.14 × 10^4^ times higher on average; p = 4.78 × 10^-6^).

### Impact of future climate on timing of outbreaks

The UK climate is expected to warm considerably over the remainder of the century^30^ and a general warming trend was clear when historic temperatures (average daily maximum temperatures by month for the period covered by the dataset of disease incidents in the wild [1991-2010]) were compared to projected temperatures for the period 2070-99 under a high emissions scenario. A warmer climate will expand the geographic area where environmental conditions (average monthly maximum daily temperatures exceeding 16°C) are likely to be suitable for severe incidents of ranavirosis. For example, the suitable geographic area in May is projected to increase by 136% by 2070 compared to the historic baseline (Fig. 4). The projected changes in UK temperatures also will extend the duration of the “disease season”, creating favourable conditions for disease during the spring and autumn as well as the summer.

**Figure 4.**
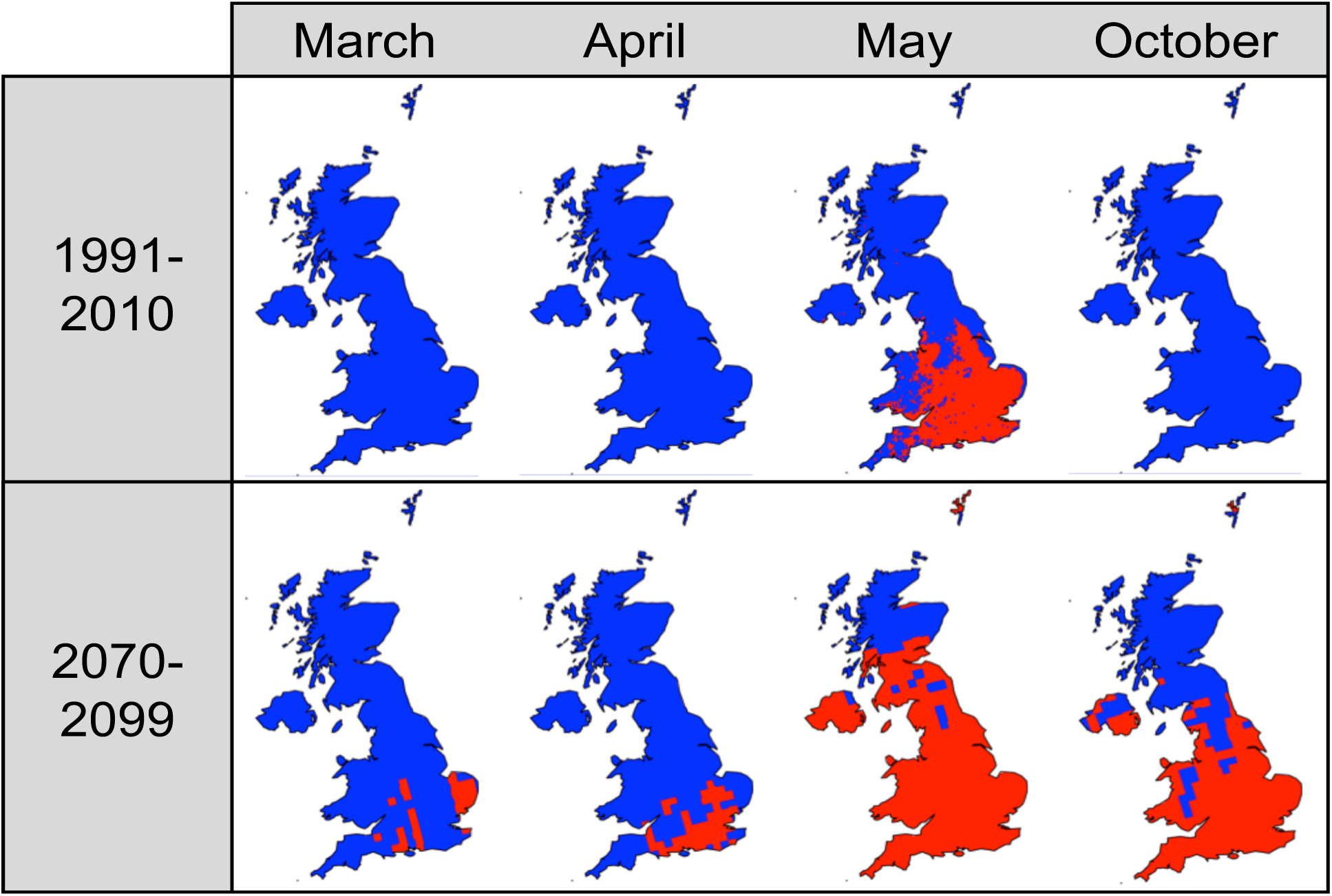
Projected shifts in geographic extent and the temporal window of ambient temperatures high enough to increase the risk and severity of ranavirus incidents in the UK. The top row of plots shows regions (red) where the average monthly temperatures in the UK (5 x 5km grid squares) exceeded 16°C for the period 1991-2010. The bottom row of plots shows regions where the projected average monthly daily maximum temperatures in the UK (25 x 25km grid squares) have a greater than 50% probability of exceeding 16°C under a high emissions scenario (IPCC scenario A1FI) for the period 2070-2099.

Temperatures during March, April and October, months which we expect to have experienced limited ranavirus incidence and severity previously, are projected to change to such a degree that this limitation will be removed for large areas of the UK (Fig. 4). The potential impacts of climate warming on disease ecology, therefore, could have critical ramifications for the continued survival of amphibian populations across the UK.

Although it is challenging to detect disease and mortality in larval amphibians, all evidence points to adult common frogs as the major life history stage and species affected by ranavirosis in the wild^23,28,31^. This observation is intriguing since larval forms are usually more affected by FV3 elsewhere in the world and common frog larvae have been shown to be highly susceptible to wild-type ranaviruses in the laboratory^32–34^. Our findings suggest that temperature could be an important determinant of this partitioning of disease among life-history stages, with common frog larvae metamorphosing before pond water reaches a temperature high enough to trigger outbreaks of ranavirosis. This situation could be altered as the climate warms and the disease season is lengthened. Shifting the timing of frog disease outbreaks will alter the life history stages at risk. If common frog tadpoles become affected, the abundance of susceptible hosts will be increased with concomitant impacts on the ranavirus basic reproductive number (R0)^11^; i.e. the dynamics of outbreaks will be fundamentally changed, making predictions of their impacts more challenging. Whilst the breeding phenology of some UK amphibian species has altered in response to climatic changes, common frog breeding has not responded to increasing absolute temperatures^35^, so any compensatory change in host behaviour may be negligible. Common frog populations in regions where temperatures become suitable for severe disease outbreaks during the larval stage might experience reduced recruitment and a subsequent reduction in their capacity to persist in the presence of infection. Additionally, altering the timing of outbreaks could create opportunities for host jumps if other potential host species have increased contact with (high levels of) virus, possibly altering ranavirus associations and speciation^36^.

### Conclusions

By combining the results of *ex situ* experiments with the analysis of two types of epidemiological datasets of disease outbreaks in wild amphibian populations, we have revealed a pattern of remarkably consistent evidence supporting a substantial effect of temperature on ranavirus disease dynamics. Temperature determines both the incidence and severity of disease outbreaks through higher temperatures increasing viral growth rate which, in turn, manifests as an increased chance of disease incidents in both experimental and wild frog populations. Our results combined with climate projections show how climate change will likely play a role in shaping future ranavirus disease dynamics in UK common frogs, altering both the geographic extent and the length of the temporal window of heightened disease risk and severity. The effects of temperature are predominantly on the virus rather than the host as evidenced by: 1) cell culture experiments which eliminate host factors that might be altered by temperature and therefore attribute changes in ranavirus growth to the pathogen itself; 2) increased virulence at higher temperatures is unlikely to be counteracted by any negative effects of higher temperature on ranavirus, as viral particles are persistent in the environment and are highly tolerant of exposure to extreme temperatures despite failing to replicate at temperatures above 30°C^23,37^, and 3) we have previously shown that expression of host immunogenetic responses to ranavirus suggest that ranaviruses are successful immune evaders^38^, thus negating any possible impacts of more productive host immune responses with raised ambient temperatures.

Our laboratory findings that virulence is reduced at lower temperatures, that frogs might be better able to manage infections at lower temperatures, and field records showing the significant effects of shading, pond volume and vegetation (which might also be due to lowered temperatures of frogs) in models of incidence and severity, point to possible steps for mitigation. Thermoregulatory behaviour leading to an increase in body temperature above normal range (“behavioural fever”) was shown to reduce the odds of infection with chytrid fungus in Panamanian golden frogs^39^ and is known to be important for disease mitigation in other ectotherms^40^. Whether “behavioural cooling” also serves as an amphibian strategy for managing infections remains to be elucidated and staying cool can be a more difficult challenge than getting warm for many ectotherms^41^. However, the provision of suitable opportunities for behavioural regulation of body temperature in the form of shading, log piles and larger ponds might help manage the severity of outbreaks.

Ranavirosis has been shown to have a major impact on common frog populations in south-east England^26^ and the current study suggests that these impacts may become greater and more widespread if future climate change projections are realized. Our results - as well as the predictions that follow from them - are strengthened through the use of a model-system that allowed us to check for patterns underlying the wild epidemiology using laboratory experiments, both at the cellular and whole animal levels, and together they present a very clear case of the environment modulating an important host-pathogen interaction.

## Methods

### Temperature as a predictor of reported frog mortality and incident severity

**Temperature-dependence of ranavirus incidence**: The FMP dataset contained 4385 records for the period 1991-2010 after removing records with errors and essential missing data. The dataset was filtered for ranavirus outbreaks based on the presence of indicative signs of disease (‘ulceration’, ‘red spots on the body’, and ‘limb necrosis/loss of digits’) and a minimum of five dead animals (i.e. using the same criteria as Cunningham^23^ and Price et al.^24^) to create a binary variable describing the ranavirus status of each incident. Data on the timing of the onset of mortality (available at the resolution of month only) and the incident location were used to download the monthly average of the daily maximum temperature for each incident from the Met Office UKCP09 dataset^42^ (further details in the Supplementary methods).

The relationship between temperature and the probability of a ranavirus-positive observation was modelled as a sigmoid (logistic) transition between an upper and lower mean frequency. The four parameters of the curve (upper and lower limits, the location and the slope of the transition) were fitted using the *mle2* function in the R package, *bbmle*^43^. Starting values for the *mle2* algorithm were obtained for slope and location (intercept), using the *glm2* function^44,45^ with a bespoke link function for binomial data (Supplementary File S1). A confidence interval around the fitted line was generated using the delta method to compute variance of a function with the *emdbook* package^46,47^. The model was compared using Akaike’s Information Criterion (AIC) to a standard logistic curve fitted by *glm2*.

**Effect of increased temperature on the severity of incidents**: In order to analyse the effect of temperature on severity, a measure that includes the total number of dead animals, the dataset was filtered using only signs of disease as criteria for differentiating between incidents caused by ranavirus and those caused by other factors. Incidents were considered ranavirus-consistent if any two of the three indicative signs of disease used above were reported. The total number of dead frogs and the estimated number of surviving frogs were combined as the response variable in a generalised linear model using the quasibinomial family to account for overdispersion in the data^48^. The average local maximum temperature for the month of onset of mortality incidents and the predicted ranavirus status, as well as the interaction between main effects, were used as predictors of severity. This basic model was also extended by incorporating these same terms as well as other variables describing the pond environment (presence of other species and physical characteristics of ponds).

### Temperature preceding ranavirus outbreaks with precise timestamps

The recent dataset on ranavirus screening of mortality incidents in UK herpetofauna using molecular methods^28^ was filtered to include only those incidents that had a precise georeference (postcodes or grid references) and timestamp (date found, submitted, or examined if a post-mortem exam was conducted on receipt, which were each assumed to approximate closely to the day of death due to the rapid decomposition of amphibian carcases). The filtered dataset contained a total of 197 incidents with 31 for which ranavirus had been detected. All incidents were overlaid on the UK grid of 5 x 5 km squares and the maximum daily temperatures for the date matching the timestamp and the 50 days prior to the death(s) were extracted from plain text data files in the UKCP09 dataset^49^, which were downloaded using the R package *Rcurl*.

Ranavirus status was used as the response variable in a series of logistic regression analyses (generalised linear models using the binomial family and the logit link function) with the average maximum daily temperature in the week preceding the mortality incident and the number of consecutive days in the previous seven where temperature exceeded 16°C as predictors. These models were also run with region (Government Office Region) or latitude as an additional predictor to further control for any effect of spatial variation in temperature. As above, seasonal variation in the detectability of amphibians was controlled for by inclusion of mortality incidents caused by factors other than ranavirosis.

## Virus growth *in vitro*

The effect of temperature on the growth of two UK isolates of FV3 (RUK11, isolated from the kidney of a diseased common frog that died with systemic haemorrhages in Middlesex in 1992, and RUK13, isolated from the skin of a diseased common frog that died with skin ulceration in Suffolk in 1995^23^) was investigated in two cell lines (epithelioma papulosum cyprini [EPC, derived from the fathead minnow fish, *Pimephales promelas*^50^; ECACC 93120820] and the iguana heart reptile cell line [IgH2; ECACC 90030804]). Cells were grown on 96 well plates until more than 90% confluent and then inoculated with virus in a ten-fold dilution series ranging from a multiplicity of infection of approximately 2 x 10^-6^ to 2 x 10^3^ (five wells per dilution with an additional one well per dilution receiving a sham exposure of cell culture media only as a negative control). Titres of viral isolate stocks were equalised by reference to qPCR scores (following Leung et al.^51^). Plates were incubated at six temperatures (10, 14, 18, 22, 26, and 30°C) and monitored daily for cytopathic effect (plaques in the cell layer). After six days, plates were scored for viral growth by counting the number of replicates at each dilution where cytopathic effect was evident and calculating the Tissue Culture 50% Infective Dose (TCID50) using the method of Reed & Muench^52^. The effects of temperature and cell line on viral growth (the mean titres of the two isolates of each type) were analysed using a linear model in R.

## *In vivo* assessment of effect of temperature on virulence

All experimental procedures and husbandry methods were approved by the ZSL Ethics Committee before any work was undertaken and procedures were performed under UK Home Office license P8897246A.

Temperature was maintained by placing five individually-housed frogs selected at random from each of the three exposure treatments into one of four climate-controlled chambers constructed from polystyrene boxes - two held at 20°C and two held at 27°C (Fig. S5; see the Supplementary methods for a comprehensive description of the set up). RUK13 was used at a titre of 1.58 x 10^5^ TCID50 mL^-1^ and RUK11 at a titre of 1.58 x 10^7^ TCID50 mL^-1^ (details of preparation of inocula and exposure methods are in the Supplementary methods). Individuals reaching humane endpoints and all surviving individuals at the end of the experiment (day eight post-exposure) were euthanized following a schedule 1 method for amphibians. The number of hours post-exposure that animals were found dead or at endpoints was recorded for survival analysis.

Following death, approximately 20mg liver samples were taken from each individual and stored at -80°C until DNA was extracted for viral load estimation. Tissues were digested overnight in batches of 20 alongside extraction negatives (reagents only, no tissue). DNA extractions were completed using DNAeasy Blood and Tissue spin columns (Qiagen) following the manufacturer’s instructions. DNA was eluted in 100µL final volumes. Viral loads were estimated using quantitative PCRs against both host and viral targets (method, reaction mix and settings followed Leung et al.^51^).

Viral load scores were subjected to a small increment by adding 10^-0^ to accommodate zero values when taking logarithms (Fig. S6). Viral load data were analysed by fitting a linear mixed-effects model to account for pseudo replication due to placement of individual experimental units inside the four climate-controlled chambers. Exposure, temperature, survival outcome and the two-way interactions were modelled as fixed effects and the climate-controlled chamber as a random effect. Survival data were analysed by fitting a mixed-effects Cox proportional hazards regression model in R using the *coxme* package^53^ with exposure and temperature as fixed effects and the climate-controlled chamber as a random effect. Model coefficients were visualised using *Forestplot*^54^.

## Effect of projected climate change on timing of ranavirus outbreaks

Baseline temperature data were generated by calculating the mean daily maximum temperature for each calendar month for the period 1991-2010 (the same period covered by our dataset of reported frog mortality incidents) for each 5 x 5km grid square used by the Met Office UKCP09 project^42^. Probabilities of mean daily maximum temperature exceeding 16°C for calendar months during the period 2070-2099 under the A1FI “high” emissions scenario of future climate change^55^ were downloaded for a grid of 25 x 25km squares at UK scale using the UK Climate Projections user interface^30^.

## Data availability

All data and code required to reproduce the analyses and figures are provided as supplementary files.

## Acknowledgements

We thank Rob Knell for help and advice about climate-chamber construction. This work was funded by NERC grants NE/M000338/1, NE/M000591/1 & NE/M00080X/1.

## Author contributions

SJP & AAC designed the *in vitro* experiments, the lab work was performed by CO & WL, and analysis was performed by SJP with help from CO. SJP designed the *in vivo* experiment with help from TWJG, the lab work was performed by SJP, WL & CS, and the results were analysed by SJP. AAC oversaw collection of the epidemiological datasets, SJP, TWJG, RAN & FB planned the analyses which were conducted by SJP with help from RAN. SJP & FB wrote the first draft of the manuscript which was edited by all authors.

